# FiTMuSiC: Leveraging structural and (co)evolutionary data for protein fitness prediction

**DOI:** 10.1101/2023.08.01.551497

**Authors:** Matsvei Tsishyn, Gabriel Cia, Pauline Hermans, Jean Kwasigroch, Marianne Rooman, Fabrizio Pucci

## Abstract

Systematically predicting the effects of mutations on protein fitness is essential for the understanding of genetic diseases. Indeed, predictions complement experimental efforts in analyzing how variants lead to dysfunctional proteins that in turn can cause diseases. Here we present our new fitness predictor, FiTMuSiC, which leverages structural, evolutionary and coevolutionary information. We show that FiTMuSiC predicts fitness with high accuracy despite the simplicity of its underlying model: it was one of the top predictors on the hydroxymethylbilane synthase (HMBS) target of the sixth round of the Critical Assessment of Genome Interpretation challenge (CAGI6). To further demonstrate FiTMuSiC’s robustness, we compared its predictions with *in vitro* activity data on HMBS, variant fitness data on human glucokinase (GCK), and variant deleteriousness data on HMBS and GCK. These analyses further confirm FiTMuSiC’s qualities and accuracy, which compare favorably with those of other predictors. Additionally, FiTMuSiC returns two scores that separately describe the functional and structural effects of the variant, thus providing mechanistic insight into why the variant leads to fitness loss or gain. We also provide an easy-to-use webserver at http://babylone.ulb.ac.be/FiTMuSiC/, which is freely available for academic use and does not require any bioinformatics expertise, which simplifies the accessibility of our tool for the entire scientific community.

## Introduction

Accurately quantifying the effect of genetic variants on the fitness of the encoded proteins is one of the open challenges in biology which, if resolved, would have a tremendous impact on the understanding and treatment of genetic diseases (1; 2; 3). The experimental approaches commonly used to quantify variant effects include different mutagenesis experiments (4; 5; 6; 7) and large-scale exome screening approaches (8; 9). However, these remain expensive and time consuming, and given the ever increasing amount of genetic data that is being generated, the number of variants of unknown significance (VUS) that are waiting to be characterized keep growing and growing (10). Moreover, the genetic bases of the majority of rare diseases are still not deciphered (11), and this is even more true for complex diseases such as cancer (12). New complementary approaches are thus needed to interpret and classify these VUS and, more generally, to gain novel insights into these matters.

In the last two decades, many computational tools have been developed to predict the phenotypic effect of genetic variants (13; 14; 15; 16; 17; 18; 19; 20; 21; 22). They are mainly based on evolutionary features combined using machine learning techniques. The most recent predictors such as (13; 21) take advantage of the advent of deep learning approaches, as enough experimental data has become available to train complex models for fitness prediction (7). These methods could in principle help accelerate the discovery of clinically relevant variants and their molecular effect, but their low accuracy and poor generalization properties are major obstacles for having a strong impact on clinical decision. In addition, black-box machine learning models do not contribute to improve our understanding of pathogenic mechanisms.

Currently, the gold standard to assess the performance of fitness prediction methods is the blind community-wide experiment called Critical Assessment of Genome Interpretation (CAGI) (23; 24), which evaluates predictors on unpublished data. CAGI allows for an unbiased assessment of the methods as well as the identification of their strengths and weaknesses. Moreover, it provides guidelines on how to translate computational predictions into clinical practice.

In this paper we present our new method, FiTMuSiC, which we used in the recent CAGI6 experiment to predict the fitness of hydroxymethylbilane synthase (HMBS) variants. We begin with a presentation and discussion of our computational approach and of its performances in CAGI6. We then showcase additional results of our method on clinically relevant variants. Our results show that FiTMuSiC achieves very good performances when applied to unseen data, which demonstrates that simple linear combination models can actually perform as well as more complex machine learning-based models.

## Methods

### Features

We briefly describe the features used by our method, which are of two kinds: structural and evolutionary. Structural features use the 3-dimensional (3D) structure of the wild-type protein as input. They include:

- Relative solvent accessibility (RSA). It is defined as the ratio (in %) between the solvent accessible surface area of a residue in its given 3D structure and in a Gly-X-Gly tripeptide extended conformation; it is computed by an in-house program (25).
- PoPMuSiC (PoP) (26). This computational tool predicts the change in protein thermodynamic stability upon point mutations (ΔΔ*G*) using the 3D structure of the target protein as input. It is based on the formalism of statistical potentials (27), with the energy values and RSA used as features in an artificial neural network.
- MAESTRO (MAE) (28). This tool also predicts the ΔΔ*G* based on the protein 3D structure. It uses contact potentials as features, as well as some biophysical properties of the mutated and wild-type residues such as hydrophobicity and isoelectric point.
- SNPMuSiC (SNP) (15). It is a predictor of variant deleteriousness based on structural and evolutionary features. Its evolutionary part is the PROVEAN algorithm (16), and its structural part consists of statistical potentials and RSA appropriately combined with artificial neural networks (ANN). We used here the structural part only, since PROVEAN is used by FiTMuSiC as a separate feature.

FiTMuSiC also includes four evolutionary features. To compute them, we generated a multiple sequence alignment (MSA) of the target sequence using JackHMMER (29) (database = UniRef90 (30), iterations = 1, E-value threshold = 0.01). The evolutionary features are:

- PROVEAN score (PVS) (16). It is a pure evolutionary tool that predicts the functional effect of variants. We used a re-implemented in-house version of the program which has some small differences with respect to the original version. Namely, the MSAs are not rectified for each possible mutation in the sequence and remain unchanged from the MSAs generated by the wild-type sequence.
- Conservation Index (CI) (31). It is calculated from *f*_*i*_(*a*) and *f* (*a*), the normalized frequencies of amino acid *a* at position *i* in the MSA and in the full MSA, respectively, which are computed as:

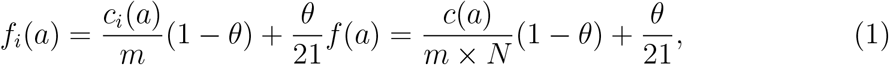

where *c*_*i*_(*a*) and *c*(*a*) are the number of occurrences of *a* at position *i* and in the full MSA, respectively, *m* is the depth of the MSA and *N* its length. The pseudocount parameter *θ* is set to 0.01 and defines the strength of the normalization; 21 is the number of possible states (20 amino acids and 1 gap). The CI score is calculated as:

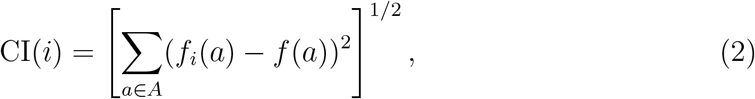

where *A* is the set of 20 standard amino acids.
- Log-odd ratio score (LOR) (32). The log-odd ratio of observing the wild-type amino acid *wt* with respect to the mutated amino acid *mt* at position *i* is defined as:

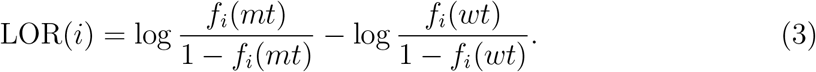
- pyCoFitness score (PYF) (33). This score is obtained through a method that infers a coevolutionary model from the MSA using a pseudo-likelihood maximization direct coupling analysis approach (34), and employs the inferred model to compute the change in fitness due to the variant.

### Model structure and training

The FiTMuSiC model is a simple linear combination of the eight features listed above. The mathematical expression of the model is:

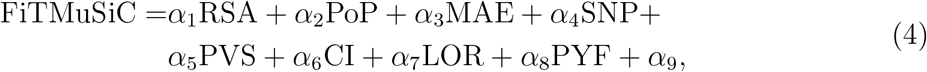

where *α*_*i*_ (*i* = 1, …, 9) are free parameters that were identified based on a training set of deep-mutagenesis scanning data on three proteins: SUMO-conjugating enzyme UBC9 (UBE2I), small ubiquitin-related modifier 1 (SUMO1) and thiamin pyrophosphokinase 1 (TPK1) (35). The structural features were computed on all training and test proteins using the AlphaFold DB structural models (36).

The conventions of FiTMuSiC values are the following: one means equal fitness for wild-type and mutant; zero means the mutant is not fit at all; values larger than one mean that the mutant is fitter than the wild-type.

### Additional models submitted to CAGI6

In addition to FiTMuSiC, we submitted the predictions of two other models to the CAGI6 challenge. The first is a simple rescaling of the SNPMuSiC score (SNP):

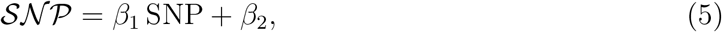

where the numerical factors *β*_1_ and *β*_2_ were chosen to rescale the SNPMuSiC values and were identified on the fitness training set described in the previous subsection.

Although stability and fitness are imperfectly correlated (37), we also submitted a prediction model based on a rescaling of the score of the thermodynamic stability predictor PoPMuSiC (POP):

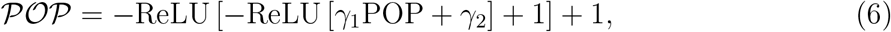

where the parameters *γ*_1_ and *γ*_2_ were identified on the same training set as the other models. The ReLU functions bound the output between 0 and 1.

### Model interpretation

To give information about the molecular effect of variants, FiTMuSiC provides four scores in addition to the global fitness of the variants. The first is the RSA of the mutated residue, which provides information on its spatial location in the 3D structure. The second is the z-score *Z* defined as:

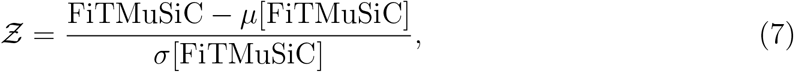

where *μ* and *σ* represent the mean and standard deviation over all mutations on the given protein, respectively. Negative z-scores correspond to mutants that are less fit than average mutants; positive z-scores indicate mutants that are fitter than average mutants, with very positive values corresponding to mutants fitter than the wild-type.

The last two scores, *Ƶ*_str_ and *Ƶ*_evo_, give information about the extent to which the structural features (SNP, POP, MAE) and evolutionary features (CI, LOR, PVS, PYF) contribute to the global fitness of the considered variant. Defining the structural (STR) and evolutionary (EVO) contributions to the fitness as:

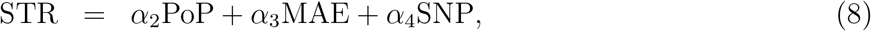

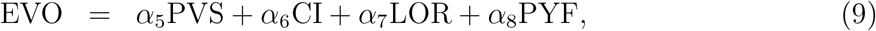

their z-scores *Ƶ*_str_ and *Ƶ*_evo_ are expressed as:

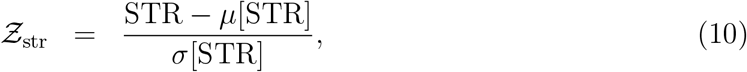

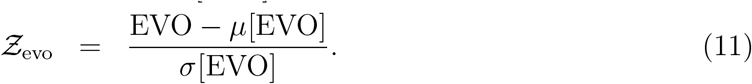

Negative *Ƶ*_str_ values correspond to mutations that destabilize the structure more than average mutations; positive *Ƶ*_str_ values indicate mutations that are less destabilizing than average mutations or are even stabilizing. Negative *Ƶ*_evo_ values correspond to mutations into residues that are rarely to never observed at that position across evolution or, more precisely, that are evolutionary unfavorable in the sequence context; positive *Ƶ*_evo_ values indicate mutations into residues that are evolutionary favorable.

## Results

### Predicting fitness of HMBS variants

HMBS, also known as porphobilinogen deaminase, is an enzyme involved in the heme biosynthesis pathway, and more specifically in the conversion of porphobilinogen into heme precursor hydroxymethylbilane (38). Mutations in this gene have been associated with acute intermittent porphyria (AIP), which is a rare metabolic disease with life-threatening neurovisceral attacks that require frequent hospitalization of patients (39). As almost one third of HMBS variants annotated in the ClinVar database (40) are VUS, saturation mutagenesis experiments using high-throughput yeast complementation assays have recently been performed to estimate the fitness of HMBS variants and better understand the pathogenic mechanisms leading to AIP (41). This data was unpublished at the time of the CAGI6 experiment and was used as blind fitness values to assess predictors.

We applied our prediction models FiTMuSiC (Eq. (4)), SNPMuSiC (Eq. (5)) and PoPMuSiC (Eq. (6)) to the HMBS target and predicted the effect of all possible single-site substitutions on its fitness. The results are reported in Table 1. We reported in the same table the results of the two other top-performing methods in the HMBS CAGI6 challenge, i.e. CalVEIR and ELAPSIC (called team 10 5 and 5 2 in the CAGI6 competition (42)), four widely used methods for deleteriousness prediction, i.e. FATHMM (43), PROVEAN (16), DEOGEN2 (14) and PolyPhen-2.0 (19), and two recently developed deep-learning based fitness predictors, EVE (13) and Sequence UNET (44). The performance of the predictors is assessed by three types of correlations between predicted and experimental fitness values of all considered variants, i.e., Pearson correlation, Spearman rank correlation and Kendall rank correlation. We also computed the root mean square deviation between experimental and predicted values (RMSD).

**Table 1:** Fitness prediction results of the benchmarked methods on the 5963 variants used in the CAGI6 HMBS challenge (42). The performances were taken from the assessors’ results, while for the other methods (below the horizontal line), we evaluated the performances ourselves. ^**\***^EVE’s predictions are available for only 5268*/*5963 variants; missing values where set to the median.

| Method | CAGI6 | Kendall | Spearman | Pearson | RMSD |
| --- | --- | --- | --- | --- | --- |
| **FiTMuSiC** | ✓ | 0.30 | 0.43 | **0.42** | **0.42** |
| *SNP* | ✓ | 0.27 | 0.39 | 0.37 | 0.46 |
| *POP* | ✓ | 0.15 | 0.22 | 0.23 | 0.46 |
| ELAPSIC team | ✓ | 0.30 | 0.42 | **0.42** | 0.45 |
| CalVEIR team | ✓ | **0.31** | **0.45** | 0.36 | 0.53 |
| FATHMM |  | 0.16 | 0.23 | 0.18 | - |
| PROVEAN |  | 0.22 | 0.32 | 0.31 | - |
| DEOGEN2 |  | 0.23 | 0.33 | 0.21 | - |
| PolyPhen-2 |  | 0.21 | 0.28 | 0.22 | - |
| EVE* |  | 0.29 | 0.42 | **0.42** | - |
| Sequence UNET |  | 0.20 | 0.29 | 0.28 | - |

FiTMuSiC performs as well as the other two best performing predictors in the CAGI16 challenge, ELAPSIC and CalVEIR (Table 1). If we take the RMSD as performance measure, FiTMuSiC outperforms all predictors. If we take the Kendall or Spearman rank correlations, CalVEIR appears to be the best. Choosing the Pearson correlation places ELAPSIC and FiTMuSiC as the best. Note that the absolute differences of the scores between the three methods is small. These three predictors all perform significantly better than those of the other teams participating in CAGI16 (42). They also perform better than all the other methods tested but one, *i*.*e*. FATHMM, PROVEAN, DEOGEN2, PolyPhen-2 and Sequence UNET. EVE is the only method that achieves performances similar to those of the three top-performing CAGI6 methods.

Note that the current version of FiTMuSiC (available on our webserver) slightly outperforms the version used for the CAGI6 HMBS challenge due to a small implementation modification. Namely, we now considered the SNP and PVS features separately (as described in Methods), whereas they were aggregated into a single term in the previous version. Indeed, the Kendall, Spearman, Pearson correlations as well as the RMSD improved from (0.30, 0.43, 0.42, 0.42) to (0.31, 0.45, 0.44, 0.41).

We also wish to underline the good performances of the SNPMuSiC deleterious variant predictor (15), which only slightly underperforms the best methods. In contrast, PoPMuSiC (26), which predicts stability changes upon mutations, does not work so well. This is not surprising given deleteriousness and fitness are very well correlated, while stability and fitness are less so. For example, all functional residues are highly important for fitness while very poorly optimized for stability (37).

The performance of the tested methods can be considered as good considering that the HMBS data was not seen by any of the methods. However, there is still room for improvement as the Pearson correlation coefficient of all methods is below 0.5. Note, however, that the noisiness of deep-mutagenesis datasets (with both random and systematic errors) puts an upper bound to the precision of the predictors which cannot be surpassed without overfitting.

### Feature analysis and model interpretation

It is well known that enzymes exhibit an activity-stability trade-off: residues in catalytic regions are optimized for functional reasons and less or not at all for stability, while other residues are very important for protein folding and stability and play little to no role in function (37). FiTMuSiC can help in distinguishing these functional and structural contributions. Indeed, it outputs the z-scores *Ƶ*_str_ and *Ƶ*_evo_ (Eqs (10-11)) which inform us about the extent to which structural and/or evolutionary features contribute to protein fitness, and provides us with a molecular-level understanding of variant effects. It also gives us information about the RSA of the mutated residues, and thus about their location in the protein.

We focused here on three functionally or structurally important residue groups of HMBS, which are structurally represented in Fig. 1 and colored according to their average per-residue z-score values *Ƶ*_evo_ and *Ƶ*_str_. Paired *Ƶ*_str_ and *Ƶ*_evo_ values of all single-site mutations are plotted in Fig. 2, with the mutations of the selected residue groups highlighted.

**Figure 1:**
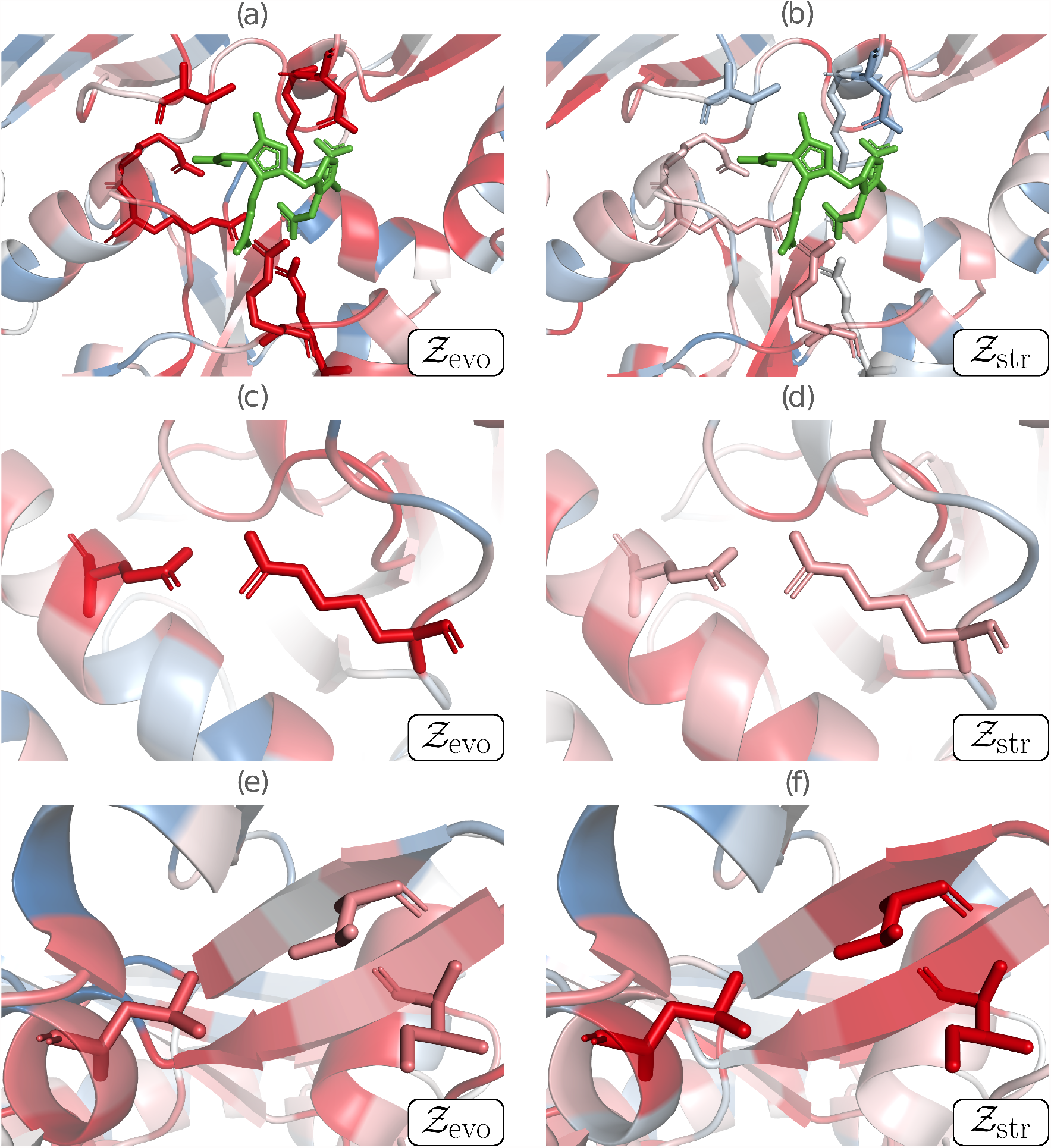
Contributions of structural and evolutionary features to HMBS fitness, represented by *Ƶ*_str_ and *Ƶ*_evo_, respectively. Negative z-scores (indicating mutations less fit than average mutations) are in red, close to zero scores in white and positive scores (indicating mutations fitter than average mutations) in blue. (a)-(b) Catalytic region, with the catalytic residues K98, D99, R149, R150, R167, R173 and C261 shown in sticks, and the substrate in green; (c)-(d) Salt bridge partners E250 and R116 shown in sticks; (e)-(f) Cluster of the three buried hydrophobic residues V124, I186 and L193 shown in sticks.

**Figure 2:**
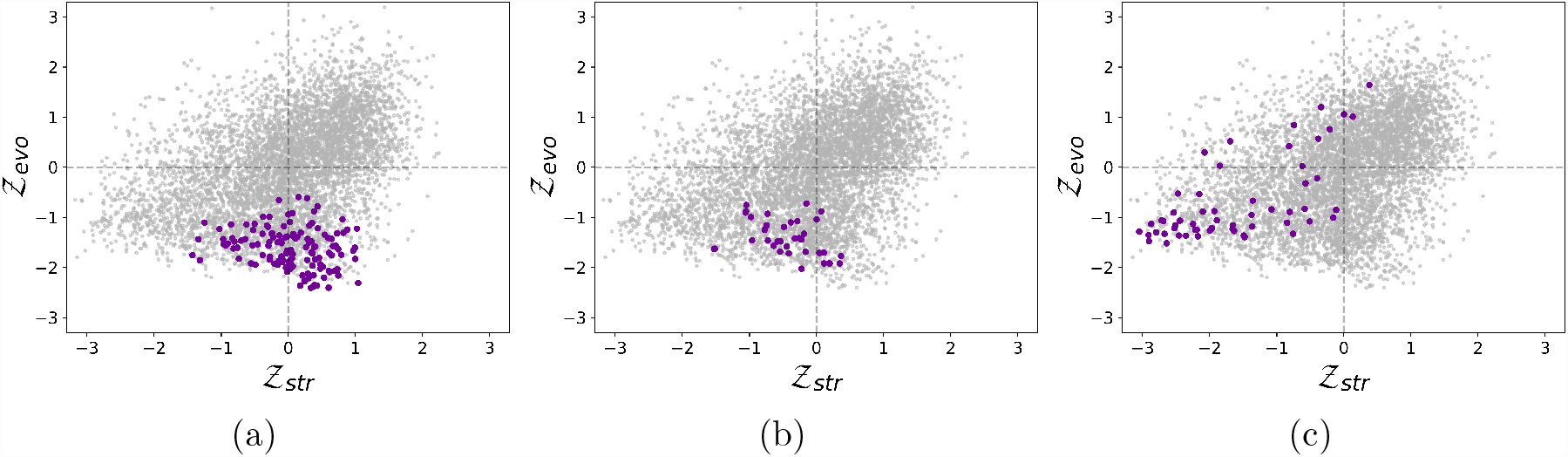
Scatter plots of paired *Ƶ*_str_ and *Ƶ*_evo_ values for all single-site mutations in HMBS. Mutations of (a) the catalytic residues K98, D99, R149, R150, R167, R173 and C261, (b) the salt bridge residues E250 and R116 and (c) the hydrophobic cluster residues V124, I186 and L193 are highlighted in purple.

The region around the catalytic site of HMBS is represented in Figs 1a-b and 2a. The catalytic residues (K98, D99, R149, R150, R167, R173 and C261) were identified by aligning the sequences of the considered human HMBS and of *Escherichia coli* HMBS, and by mapping the seven catalytic residues of the latter (45) annotated in the Catalytic Site Atlas (46). These residues are thus functionally important, well conserved and very specific. As expected, mutating them results in very negative *Ƶ*_evo_ values (between −2.41 and −0.59), which reflects drastic reduction or loss of function. In contrast, they contribute little to structural stability, as seen from the predicted *Ƶ*_str_ values centered around zero (between −1.43 and +1.04).

The second region considered is the salt bridge between the negatively charged residue E250 and the positively charged residue R116 (Figs 1c-d and 2b). It is a highly specific interaction that has been shown to play an essential role in the enzyme’s fold by molecular dynamics simulations (41). The *Ƶ*_evo_ and *Ƶ*_str_ values of these two residues are predicted to be negative on the average (−1.41 and −0.43 respectively), indicating fitness reduction upon mutations. *Ƶ*_evo_ is negative for all mutations (≤ −0.72), whereas *Ƶ*_str_ is only negative on the average (between −1.53 and +0.37). The high specificity of the interaction gives a particularly strong evolutionary signal, whereas the stabilizing effect of salt bridges is less marked compared to other interactions.

Finally, the hydrophobic cluster of the three residues V124, I186 and L193 (Figs 1e-f and 2c) located in the core of the protein is very important for the stability of the protein fold. It thus shows strongly negative *Ƶ*_str_ values, with some exceptions that correspond to mutations from one hydrophobic residue into another. In contrast, this cluster plays no direct role in the protein’s enzymatic activity and, moreover, hydrophobic interactions have low specificity and are often substituted with other hydrophobic residues across evolution. This explains the large width of the *Ƶ*_evo_ distribution (between −1.52 and +1.64), and its only weakly negative average value (−0.70). On the other hand, *Ƶ*_str_ values are also sparse (between −3.05 and +0.38) but are more shifted towards negative values (average of −1.60).

Since evolution and structure are related, it is no surprise that we often observe correlated *Ƶ*_str_ and *Ƶ*_evo_ values. However, this correlation is limited (Pearson correlation of 0.40). As a matter of fact, there are a lot of counterexamples where *Ƶ*_str_ and *Ƶ*_evo_ have opposite signs, as seen in Fig. 2. This reflects the fact that the evolutionary and structural components of fitness are complementary, and that combining them into a single model increases both its accuracy and interpretability.

### FiTMuSiC application to HMBS variant pathogenicity and activity

Fitness predictors are expected to play a crucial role in the classification and interpretation of genetic variants by providing complementary information to the experimental characterizations (47). It has however to be noted that the experimental HMBS fitness values of the CAGI6 challenge come from a deep mutagenesis experiment that uses functional complementation yeast assays, which cannot fully reflect the complex mechanisms underlying variants’ pathogenicity and activity.

In this context, we assessed all the predictors considered as well as the experimental yeast assay data (41) on their ability to distinguish clinically annotated pathogenic and benign variants in humans. To that end, we collected 53 pathogenic or likely pathogenic variants in HMBS that are related to AIP and 13 benign or likely benign variants from ClinVar (40). The metrics we used to assess the methods’ performances are sensitivity, specificity and balanced accuracy (BACC), for which we used the prediction thresholds provided by the predictors’ authors. As threshold-independent metrics, we chose AUC-ROC, the area under the receiver operating characteristic curve. We reported all performances in Table 2. We observe that FiTMuSiC predicts with very high accuracy the pathogenicity of the variants with a BACC of 0.94 and an AUC-ROC of 0.98. It performs better than all other computational methods and also, notably, than the experimental high-throughput fitness data obtained by yeast complementation assays to evaluate variant pathogenicity. We found that some of the computational methods tested are heavily biased towards pathogenic variants, as for example PolyPhen-2 and FATHMM. This can be explained by the choice of the threshold values proposed by their authors. They have thus a very poor specificity and predict very few neutral variants. FiTMuSiC does not suffer from this bias and reaches almost perfect accuracy in identifying neutral variants. Note that EVE also shows good performances which are only slightly less accurate than FiTMuSiC.

**Table 2:** Performance on 66 HMBS variants with clear clinical annotations (41), using all predictors assessed as well as experimental fitness data obtained by yeast complementation assays (41).

| Method | Sensitivity | Specificity | BACC | AUC-ROC |
| --- | --- | --- | --- | --- |
| Experimental | 0.81 | **0.92** | 0.87 | 0.92 |
| **FiTMuSiC** | 0.96 | **0.92** | **0.94** | **0.98** |
| FATHMM | **1.00** | 0.00 | 0.50 | 0.79 |
| PROVEAN | 0.96 | 0.77 | 0.87 | 0.87 |
| DEOGEN2 | **1.00** | 0.23 | 0.62 | 0.93 |
| PolyPhen-2 | 0.98 | 0.31 | 0.64 | 0.91 |
| EVE | 0.94 | **0.92** | 0.93 | **0.98** |
| Sequence UNET | 0.70 | 0.77 | 0.73 | 0.82 |

As an additional verification of FiTMuSiC robustness, we checked if it is able to predict the effect of variants on HMBS *in vitro* activity. We reported in Table 3 the correlations between the results of the predictors or high-throughput experiments and the experimentally measured activity of variants (48). These results show that FiTMuSiC performs very well with the second best Pearson correlation coefficient just after EVE, and the second best Spearman rank correlation coefficient after high-throughput experimental fitness data (41).

**Table 3:** Correlation coefficients between experimental activity on 35 HMBS variants measured in (48) and the fitness values obtained by the assessed predictors and by experimental yeast complementation assays (41).

| Method | Spearman | Pearson |
| --- | --- | --- |
| Experimental | **0.77** | 0.72 |
| **FiTMuSiC** | 0.53 | 0.85 |
| FATHMM | 0.53 | 0.57 |
| PROVEAN | 0.19 | 0.50 |
| DEOGEN2 | 0.41 | 0.57 |
| PolyPhen-2 | 0.38 | 0.53 |
| EVE | 0.46 | **0.88** |
| Sequence UNET | 0.08 | 0.35 |

### FiTMuSiC application to human glucokinase

To further test the robustness of FiTMuSiC, we applied it to another blind test set containing experimental high-throughput fitness data of all single-site variants in human glucokinase (GCK). This enzyme plays a key role in insulin secretion in pancreatic *β*-cells: it catalyzes the first step of the glycolysis by transforming glucose into glucose-6-phosphate (49). Inactivating GCK variants were related to maturity-onset diabetes of the young as well as to permanent neonatal diabetes mellitus (49; 50). Hyperactive GCK variants are also deleterious and lead to persistent hyperinsulinemic hypoglycemia of infancy.

In order to shed light on the molecular effects that lead to these disorders, the GCK activity of all possible single-site variants have been experimental assessed using functional complementation yeast assays (51). We used this set of 6862 variants as independent test set to assess the fitness predictors. To follow the same benchmark conditions as for HMBS in CAGI6, we floored all negative fitness values to zero and excluded all values of which the standard error exceeds 0.3. We reported in Table 4 the performances of FiTMuSiC and other computational tools.

**Table 4:** Correlations between fitness values obtained by high-throughput experiments using functional complementation yeast assays (51) on 6862 GCK variants and those predicted by all the methods assessed. ^*^EVE’s predictions are available for only 6414 out of the 6862 GCK variants.

| Method | Spearman | Pearson |
| --- | --- | --- |
| **FiTMuSiC** | 0.49 | **0.40** |
| FATHMM | 0.39 | 0.30 |
| PROVEAN | 0.41 | 0.33 |
| DEOGEN2 | **0.51** | 0.35 |
| PolyPhen-2 | 0.36 | 0.23 |
| EVE* | 0.47 | **0.40** |
| Sequence UNET | 0.30 | 0.24 |

We found that FiTMuSiC reaches performances on this test set that are close or slightly better than the top-performing methods while having a simpler model. Indeed, FiTMuSiC has the highest Pearson correlation coefficient *ex aequo* with EVE (0.40), and the second best Spearman correlation coefficient (0.49) just after DEOGEN2.

We also evaluated the ability of the methods to classify deleterious and benign variants that are defined based on clinical annotations. For that purpose, we curated a set of variants in GCK from ClinVar (40) with clear clinical interpretation. This led us to a collection 70 pathogenic or likely pathogenic variants, and 3 benign or likely benign variants. The very low number of benign variants and the bias of predictors towards deleterious variants make this test case relatively easy, and most methods thus reach very high scores: two methods have a BACC of 0.96 or more, and four methods have an AUC-AUROC of at least 0.98 (Table 5). FiTMuSiC is also performing very well with a BACC of 0.89 and an AUC-ROC of 0.99. Due to the strong imbalance of this test set, we suggest to consider these results with caution.

**Table 5:**
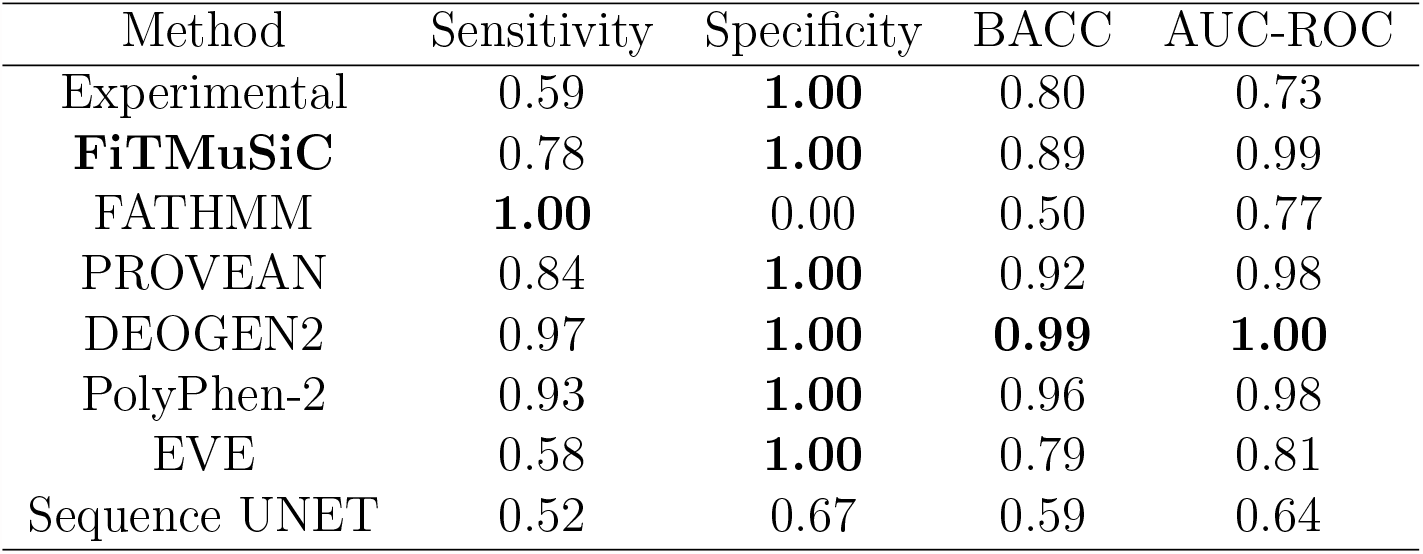
Performance on 73 GCK variants with clear clinical annotations taken from ClinVar (40), using all predictors assessed as well as experimental fitness data obtained by functional yeast complementation assays (51).

It has to be noted that the use of experimental fitness data from complementation yeast assays to predict deleteriousness does not perform very well for GCK variants (Table 5). The BACC and AUC-ROC values are even lower than in the case of HMBS. Some reported pathogenic variants such as V62M, T65I and H137R, seem to be benign in the experimental fitness map. Their deleteriousness has been suggested to be related to effects such as modest structural instability which are not captured by the assay (51).

### Webserver

In order to make FiTMuSiC readily available to the scientific community, we have developed an easy-to-use webserver at http://babylone.ulb.ac.be/FiTMuSiC/. Users need to input a 3D structure of the target protein in one of three ways:

1. Provide its PDB ID if it is available in the Protein Data Bank (PDB) (52); the structure is automatically retrieved.
2. Provide its UniProt ID; the corresponding AlphaFold DB structure (36) is then retrieved.
3. Provide a personal structure in PDB format (.pdb).

Since FiTMuSiC provides results on a per-chain basis, users need to select which chain they want the results for. Note that FiTMuSiC only outputs the results of a single chain, but the structural components of the model take into account all the chains contained in the structure file when computing the fitness score. Therefore, we recommend that users provide protein structures that correspond to biological units, especially when dealing with multimers.

Once the chain has been selected and submitted, the computation starts. Depending on the length of the query protein and the depth of its MSA, users should expect the computation to be completed in a few minutes for short proteins to a few hours for very long proteins. Once the computation is done, a CSV file with the results is sent to the email address provided during the submission. This file contains the RSA of all residues in the protein and the predicted fitness scores for all possible single-site variants. The last four columns contain fitness score information, i.e. the raw FiTMuSiC score and the z-scores *Ƶ, Ƶ*_evo_ and *Ƶ*_str_ (Eqs (4),(7),(10),(11)). When the score of a given residue mutation is computed in the absence of structural information for that residue, or based on an MSA of very small depth, a warning is attached to inform about the possibly lower accuracy of the prediction.

More information about the webserver and its usage is available on the help page (http://babylone.ulb.ac.be/FiTMuSiC/help.php).

## Conclusion

We presented here FiTMuSiC, our new computational model based on a combination of structural and (co)evolutionary information, which predicts the impact of single-site amino acid substitutions on protein fitness. We applied it to predict variants in HMBS, one of the targets of the CAGI6 challenge. It was rated as one of the top three predictors by the CAGI6 assessors (42). The strengths of FiTMuSiC can be summarized as follows:

- It is based on a simple model, which is less prone to overfitting and biases towards the training set than machine learning models with thousands of parameters. This allows for very good performances on blind, independent test sets as we have shown here for variants in HMBS and GCK.
- It retains interpretability by providing the *Ƶ*_evo_ and *Ƶ*_str_ scores which allow distinguishing between variants that impact more on function or on stability.
- It is available through an easy-to-use webserver, which allows users to get FiTMuSiC results in a simple way even without bioinformatics background.

For all these reasons, FiTMuSiC is of interest to the large community of scientists interested in the prioritization, classification and interpretation of genetic variants. Moreover, it represents a reliable, complementary and cheaper approach compared to experimental methods.

## Acknowledgements

We would like to thank the assessors of the CAGI6 HMBS challenge Jing Zhang, Qian Cong and Nick V. Grishin for their assessment work and for providing us detailed information about it. We thank Steven Brenner, Predrag Radivojac and Constantina Bakolitsa for organizing the CAGI competition as well as the experimental group led by Frederick P. Roth for providing the data on HMBS variant fitness used as blind test in the HMBS challenge.

